# Dogma and belief: the primary lesion in age-related hearing loss is old news

**DOI:** 10.1101/2022.01.14.472488

**Authors:** David McAlpine, Heivet Hernandez-Perez, Mark Seeto, Brent Edwards

## Abstract

Presbycusis, or age-related hearing loss, is the most common sensory deficit globally, and the biggest modifiable risk factor for a later dementia diagnosis. Despite its ubiquity, however, the primary pathology contributing to presbycusis is reportedly contentious, particularly the relative role of damage to the sensory outer hair cells compared to the stria vascularis, an important inner ear structure that maintains the ionic concentration of inner ear fluids that surround it. To determine what might be the “dogma” of the field regarding the primary pathology in presbycusis, we conducted an online Google survey (https://forms.gle/GPreoePmRxBBkchc7) asking relevant respondents in the field their opinions on the matter. In Question (Q1), respondents were asked to rate ‘*in your opinion*’ from ‘least likely’ to ‘most likely’ (on a scale 1 to 4 [being ‘most likely’]) ‘*what is the primary pathology contributing to presbycusis?*’ in terms of ‘*damage to*’: i) the inner hair cells, ii) outer hair cells (OHCs), iii) spiral ganglion, iv) stria vascularis. The term ‘dogma’ suggests that the proportion of people in the field who believe that the main cause of presbycusis is damage to the stria vascularis is at least 50%. The results of our survey estimated this proportion to be 19/101 = 0.188 (95% CI [0.124 0.275]) and a two-sided test of the null hypothesis that this proportion is at least 0.5 was rejected (*p* < 10^−10^). When it came to opining what ‘*other professionals in the field consider to be the primary pathology contributing to presbycusis*’ (Q2), the tendency to rank ‘damage to OHCs’ as being the primary pathology was 45%. S*tria vascularis* was least likely to be ranked 4 (11%) by professionals in the field opining about the beliefs of others. Even when *‘noise damage was excluded’* as a factor (Q3), the ‘most likely’ contributing factor to presbycusis was stated to be damage to the OHCs (42%). Our data suggest the dogma of the field is that damage to outer hair cells is the primary pathology in presbycusis.

## Introduction

Presbycusis, or age-related hearing loss, is the most common sensory deficit globally (Tu & Friedman, 2018; Wattamwar et al., 2017; World Health Organization, 2021), and the single biggest modifiable risk factor for a later dementia diagnosis (Lin et al., 2013; Livingston et al., 2017). Despite its ubiquity, however, the primary pathology contributing to presbycusis is reportedly contentious (Lee, 2013; Wu et al., 2020; Yu et al., 2021). Demonstrating evidence consistent with presbycusis as primarily an outcome of damage to sensory outer hair cells of the inner ear, Wu and colleagues concluded their findings overturned the dogma in the field, namely that the primary lesion underlying presbycusis is believed to be damage to stria vascularis, the structure within the inner ear that generates and maintains the ‘battery’ in hearing, the endocochlear potential, essential to mechano-electrical (and electro-mechanical) transduction by sensory hair cells. That these findings overturn the ‘dogma’ in the field was surprising to us, as our own beliefs are that OHCs do indeed represent the primary lesion in presbycusis, and it is our opinion that colleagues in the general field of hearing and deafness generally concur with this view. Thus, consistent with *the “dogma of the field”*—the primary definition of ‘dogma’ in the Merriam-Webster Dictionary being ‘*something held as an established opinion*’—we sought to determine what might be the established opinion held by self-recognised persons in the field of hearing sciences and healthcare (scientists, technologists, clinicians, and the like) as to the primary site of pathology in presbycusis. Given current global efforts to restore hearing function in adults with age-related hearing loss, it is important to understand what those seeking to abolish, reverse, or ameliorate the effects of presbycusis believe to be the most relevant reason for hearing loss acquired over the life course. Using an online survey targeted at relevant persons, we find that the majority opinion is strongly biased towards outer hair cells being the primary lesion site in presbycusis, that this opinion is even more strongly held when respondents are asked to consider what others might believe to be the case, and that this still holds when exposure to loud sounds—a known contributing factor to hair cell damage—is excluded as a potential contributing factor. Our data provide support for the view that the dogma of the field is that damage to outer hair cells is the primary lesion in presbycusis.

## Methods

We conducted an online (Google) survey asking potentially relevant respondents in the field their opinions as to the primary pathology in presbycusis. This research met the requirements set out in the National Statement on Ethical Conduct in Human Research (2007) and was approved by Macquarie University’s Medicine & Health Sciences Subcommittee (Reference No: 52020795021789).

A unique link to a Google survey (https://forms.gle/GPreoePmRxBBkchc7) was advertised via email lists and social media in the broad domain of hearing and deafness. Once the questionnaire was accessed, potential respondents were informed that the survey inquired about the current belief of researchers, clinicians, and technologists in the field of hearing and deafness as to the primary pathology contributing to presbycusis (age-related hearing loss) as well as potential therapeutic solutions. To complete the survey, respondents were required to self-identify as a researcher, educator, clinician, clinician-researcher, or technologist in the field of hearing and deafness and being 18 years old or above. The survey comprised eleven questions in total: (1-4 related to participants’ age, profession, and region where their research and/or work was conducted). Questions 5-7 corresponded to specific participants’ beliefs of the causes of age-related hearing loss whilst the remaining 4 questions corresponded to participants’ views on therapeutic solutions. We considered only questions related to participants’ beliefs of the causes of age-related hearing loss (questions 5-7, hereafter referred to as Q1-Q3).

The distribution of responses for each combination of role and damage location were calculated for Q1-Q3. The specific null hypothesis of the stria vascularis being the most important cause of age-related hearing loss was tested. If the null hypothesis is supported, then the statement that the dogma of the field is that damage to the stria vascularis is the primary cause of presbycusis would be refuted. In addition, Spearman correlations between the responses for each location of damage in Q1 and the responses for each location of damage in Q2 and Q3 were also analysed.

## Results

### Beliefs on the primary pathologies contributing to presbycusis

After securing basic demographic data from the 101 respondents, including self-identified role—clinician (31), educator (5), researcher (43), researcher-clinician (15), technologist (7)—we asked respondents three questions concerning their opinions as to the primary pathology in presbycusis. In Question (Q1), respondents were asked to rate ‘*in your opinion*’ from ‘least likely’ to ‘most likely’ (on a scale 1 to 4, with 4 being ‘most likely’) ‘*what is the primary pathology contributing to presbycusis (age-related hearing loss)?*’ for each of four structures in the inner ear, in terms of ‘*damage to*’: i) the inner hair cells (IHC), ii) outer hair cells (OHC), iii) spiral ganglion (SG) iv) stria vascularis (SV). The data indicate a preference for the belief that damage to the outer hairs is the primary pathology/site of action for presbycusis. The highest ranking of 4—’most likely’—returned the highest proportion of responses (38%), and two thirds of respondents (68%) responded with either 3 or 4, the two highest rankings, for outer hair cells being the primary pathology, with just 20 % of respondents suggesting the lowest ranking of 1—least likely—for ‘damage to outer hair cells’.

In terms of damage to stria vascularis, the greatest number (30%) of respondents responded with ranking 2, and the smallest (19%) ranking 4—’most likely’, with a slight majority (52%) of respondents responding with the combined lowest rankings 1 (‘least likely’) and 2 for stria vascularis being the primary site of damage in presbycusis. ‘Damage to the spiral ganglion’ garnered least support, with nearly 70% of respondents scoring it either 1 or 2 (i.e., low). Respondents were more likely to rank damage to inner hair cells as being the contributing factor in presbycusis, with ranking 4 (‘most likely’) accounting for the largest number of respondents (28%) and the more/most likely rankings 3 and 4 combined for a total of 55% of respondents. Overall, for the four potential sites of action listed in our survey, outer hair cells were considered most likely, and stria vascularis least likely, (see Figure 1 for details).

**Figure 1.**
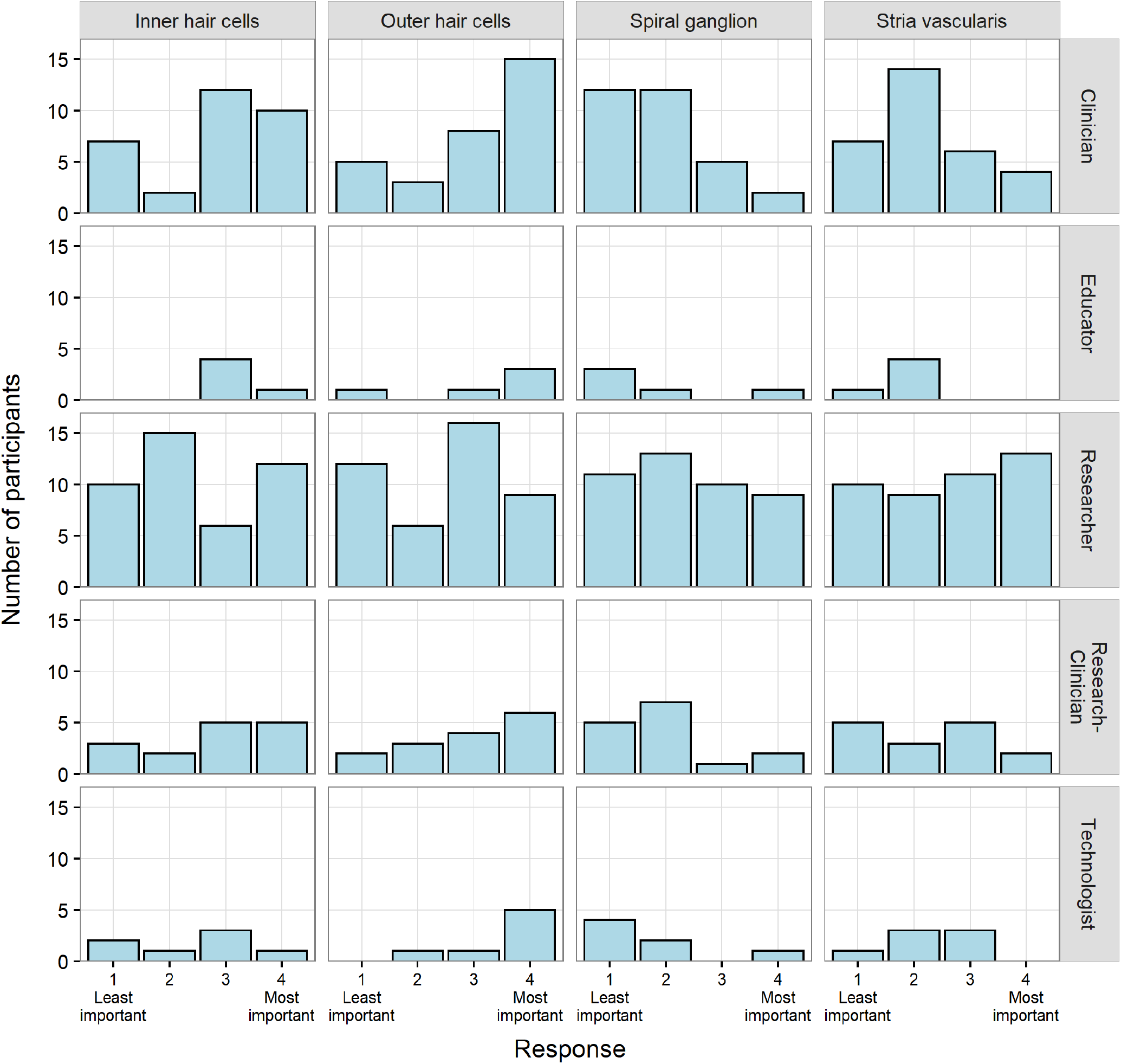
Distribution of responses for Q1 (what you believe). The underlying data can be found in https://doi.org/10.25949/17009321.v1

The term ‘dogma’ suggests that the proportion of people in the field who believe that the main cause of presbycusis is damage to the stria vascularis is 50% at the very least. Based on our sample of 101 respondents, this proportion was estimated to be 19/101 = 0.188 (95% confidence interval [0.124, 0.275]), and a two-sided test of the null hypothesis that this proportion is 0.5 was rejected with a p-value of about 10^−10^. For a higher null value of the proportion, the p-value would be even smaller. Even considering only the group self-identified as ‘researcher’, which had the highest proportion of participants choosing stria vascularis as most important in Q1, there was still substantial evidence against the null hypothesis that the true proportion is 0.5 in this group, with a sample proportion of 13/43 = 0.302 (95% CI [0.186 0.451]), and a two-sided p value = 0.01.

When it came to opining what ‘*other professionals in the field (inc. researchers, clinicians, technologists) consider to be the primary pathology contributing to presbycusis*’ (Figure 2), the tendency to rank ‘damage to outer hair cells’ as being the primary pathology was even greater than the tendency to rank it as such in terms of one’s own opinion, with 45% of respondents ranking it 4—’most likely’, and 20% as 3. In terms of stria vascularis, respondents were of the opinion, based on their rankings, that other professionals did not rank ‘damage to the stria vascularis’ highly, with just 11% selecting 4 (most likely) and a further 18% ranking it 3. Across all our listed sites of possible pathology in presbycusis, *stria vascularis* was least likely to be ranked 4 by professionals in the field opining about the beliefs of other professionals in the field. The combined proportion of respondents across the lowest two rankings (1 and 2) was greatest for ‘damage to the stria vascularis’, with the inner hair cells (32%) and the outer hair cells (32%) equally represented in these lowest rankings. These data suggest that the beliefs of professionals in the field *about the beliefs held by other professionals in the field*, favours our own understanding that the dogma is indeed that ‘*damage to the outer hair cells*’ is the likely primary pathology in presbycusis, and ‘*damage to the stria vascularis*’ is the least likely. We note these beliefs expressed by others might represent dogma in their own right—we did not request, nor are we aware of any assessment of the beliefs of other professionals by our respondents when making their judgments.

**Figure 2.**
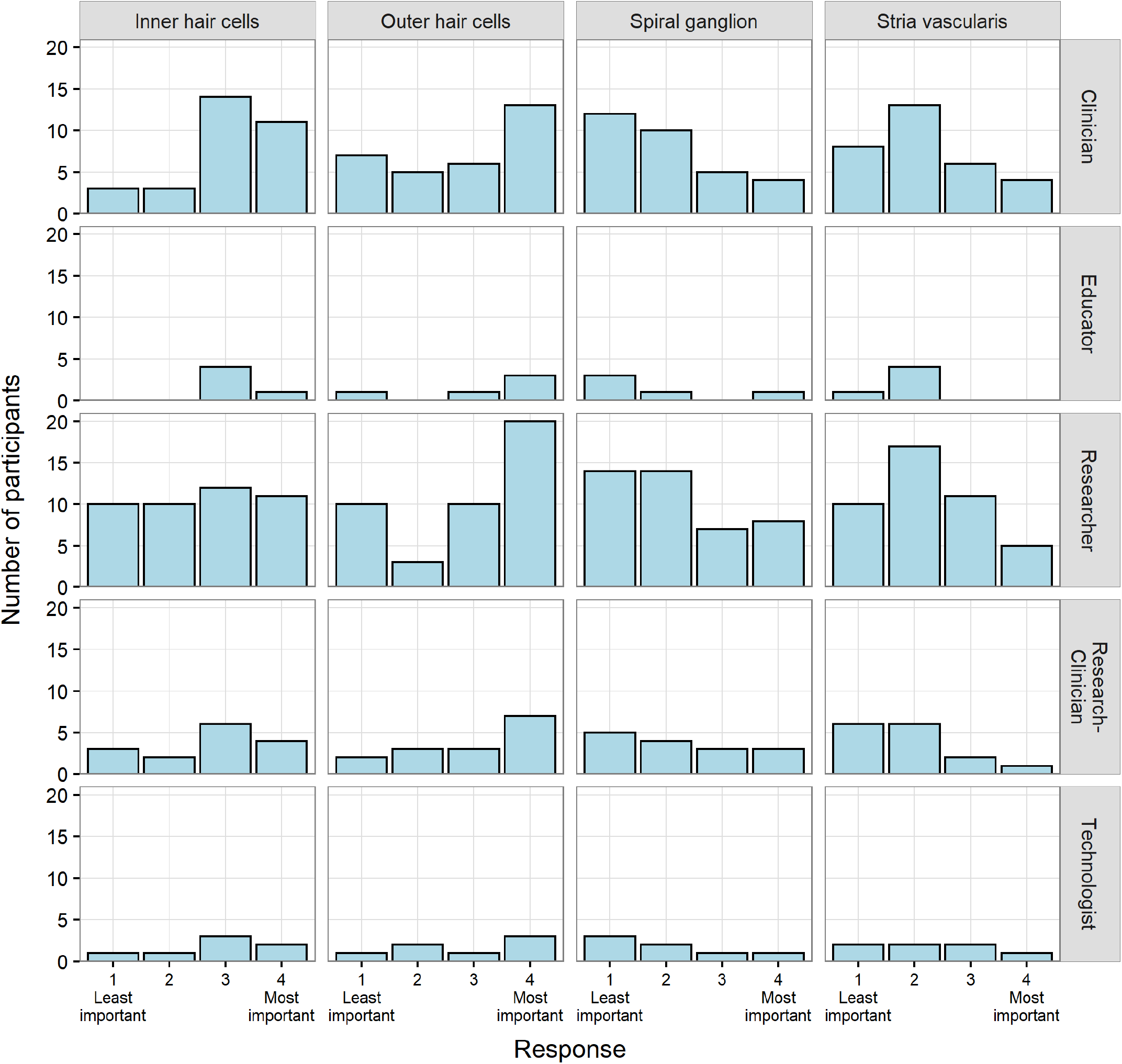
Distribution of responses for Q2 (what you think others believe). The underlying data can be found in https://doi.org/10.25949/17009321.v1

### Beliefs on the pathology underlying presbycusis when damage by loud sounds is excluded as a factor

Merriam-Webster defines presbycusis as ‘*a lessening of hearing acuteness resulting from degenerative changes in the ear that occur especially in old age*.’, succinct although ill-defined with respect to any specific site of pathology. Compelling evidence from animal studies(Kujawa & Liberman, 2009) suggests spiral ganglion neurons are a potential site of primary lesions evoked by exposure to loud sounds, generating listening problems not evident in the standard test of hearing acuity, the audiogram (which measures sensitivity to very soft sounds in quiet). Knowing the likely importance and spreading knowledge of this ‘hidden hearing loss’ (Schaette & McAlpine, 2011) in the field, we asked our respondents to rank the likelihood that each of the same four sites of potential pathology—inner and outer hair cells, stria vascularis, and spiral ganglion neurons—were the primary site of pathology once ‘*exposure to loud sounds (‘noise damage’)*’ was excluded as a possible generator of pathology (Figure 3).

**Figure 3.**
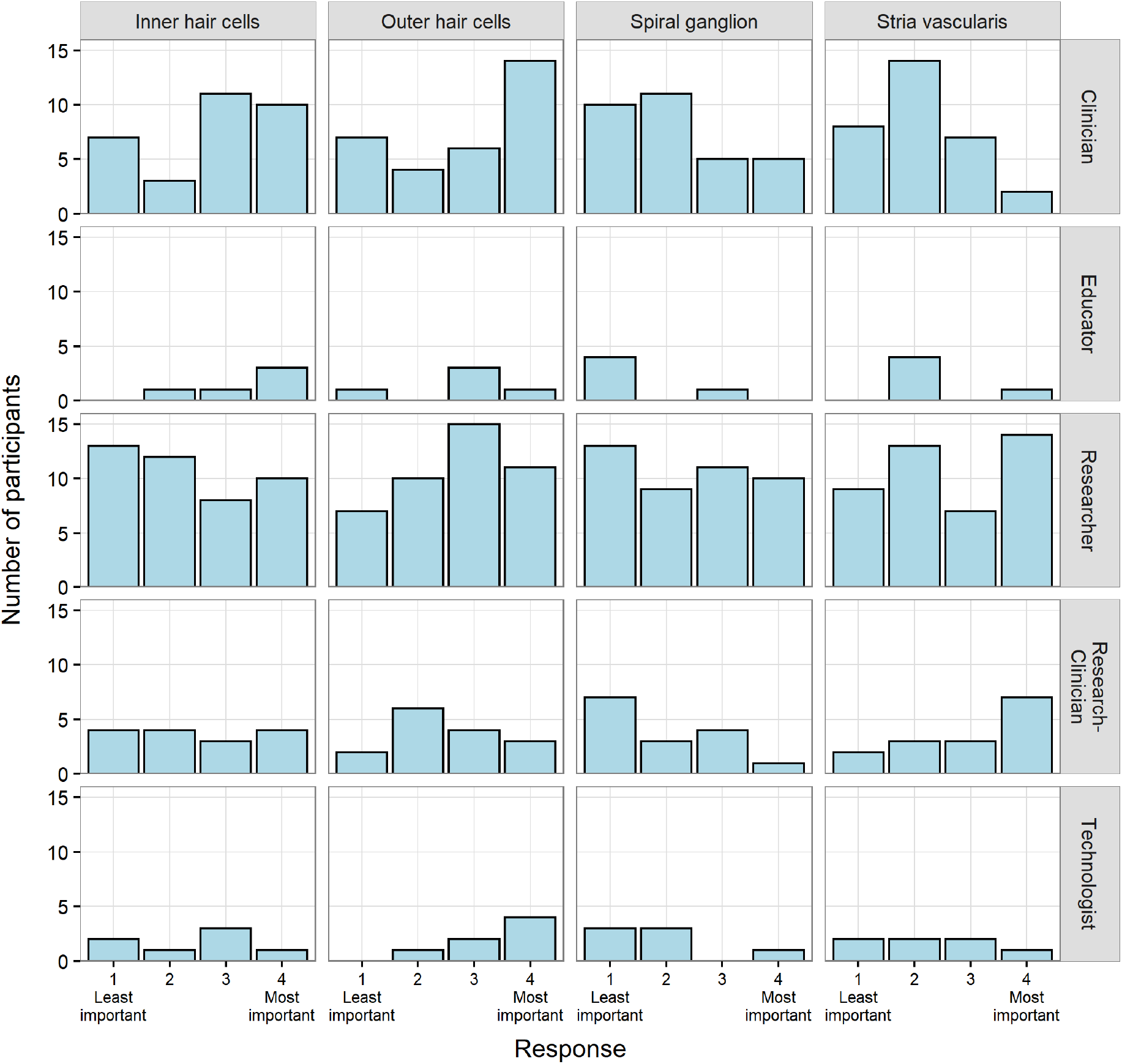
Distribution of responses for Q3 (what you believe excluding noise damage). The underlying data can be found in https://doi.org/10.25949/17009321.v1

Once more, even when noise damage was excluded as a factor, the highest ranking of 4, ‘most likely’, was attributed to ‘damage to the outer hair cells’, although this time only 28% of respondents made this judgement, against 42% when this distinction—excluding noise damage—was not made. A ranking of 4 for ‘damage to the stria vascularis’ increased from 11% to 25% upon this exclusion. Combined across rankings 3 and 4, however, it was still the case that the majority (58%) of respondents thought outer hair cells to be the likely primary site of pathology in presbycusis, compared to 42% combined across rankings 3 and 4 for stria vascularis. Excluding noise damage, only 25% of respondents considered stria vascularis as the ‘most likely’ site of pathology in presbycusis, and almost half of respondents still ranked stria vascularis as least/less likely (48% combined across ranks 1 and 2). We did not ask, but are of the opinion, that respondents’ opinions as to the opinions of others on this matter would have again favoured more strongly outer hair cells as being the primary site of pathology in presbycusis, even when exposure to loud sounds as a generator of damage is excluded. This raises a generally interesting question as to why the authors consider the dogma of the field to be such, as our data suggest professionals in the field tend to consider other professionals in the field more likely to implicate outer hair cells in presbycusis than they do themselves. The data are consistent with the dogma of the field being that damage to OHCs is the primary lesion in presbycusis whether noise damage is excluded.

It is also interesting to note that respondents were less likely, but only moderately so, to consider damage to the spiral ganglion as the primary pathology in presbycusis once noise damage was excluded; the least/less likely ranks of 1 and 2 combined increasing from 55% to 65%. This suggests that most respondents understood the term ‘presbycusis’ explicitly in terms of elevated hearing thresholds (a definition we did not prompt) and supports the view that knowledge of the spiral ganglion as a possible primary lesion site in listening problems is increasing. It is also consistent with some respondents being prompted to remember the role of noise in primary damage to the spiral ganglion—in the absence of elevated thresholds—by the order of the questions in our survey.

### Correlations between one’s own beliefs and the beliefs of others as to the primary site of pathology in presbycusis

As respondents were given the option of ranking 1-4 their own, and others’, beliefs regarding the primary site of pathology in presbycusis, we generated an average response for each question (Table 1 what you believe’; ‘what others believe’; ‘excluding noise damage’) and tested the differences in beliefs across role groups using a permutation test approach, with the null hypothesis being that the true average responses for each of the four damage locations are the same across roles. We performed this analysis for all roles combined and for the two largest role groups combined, clinicians and researchers. The p-values indicate that only for Q1 (‘what you believe’) for clinicians and researchers was the null hypothesis rejected (Q1 [p=0.03]; Q2 [p=0.31]; Q3[p=0.11]). This suggests that these two groups favour the believe that OHC are the main cause of age-related hearing loss.

**Table 1.** Average response values for Q1-Q3.

| **Q1 (WHAT YOU BELIEVE)** |  |  |  |  |
| --- | --- | --- | --- | --- |
| **ROLE** | **IHC** | **OHC** | **SG** | **SV** |
| **CLINICIAN** | 2.81 | 3.06 | 1.90 | 2.23 |
| **EDUCATOR** | 3.20 | 3.20 | 1.80 | 1.80 |
| **RESEARCHER** | 2.47 | 2.51 | 2.40 | 2.63 |
| **RESEARCHER-CLINICIAN** | 2.80 | 2.93 | 2.00 | 2.27 |
| **TECHNOLOGIST** | 2.43 | 3.57 | 1.71 | 2.29 |
| **Q2 (WHAT YOU THINK OTHERS BELIEVE)** |  |  |  |  |
| **ROLE** | **IHC** | **OHC** | **SG** | **SV** |
| **CLINICIAN** | 3.06 | 2.81 | 2.03 | 2.19 |
| **EDUCATOR** | 3.20 | 3.20 | 1.80 | 1.80 |
| **RESEARCHER** | 2.56 | 2.93 | 2.21 | 2.26 |
| **RESEARCHER-CLINICIAN** | 2.73 | 3.00 | 2.27 | 1.87 |
| **TECHNOLOGIST** | 2.86 | 2.86 | 2.00 | 2.29 |
| **Q3 (WHAT YOU BELIEVE, EXCLUDING NOISE DAMAGE)** |  |  |  |  |
| **ROLE** | **IHC** | **OHC** | **SG** | **SV** |
| **CLINICIAN** | 2.77 | 2.87 | 2.16 | 2.10 |
| **EDUCATOR** | 3.40 | 2.80 | 1.40 | 2.40 |
| **RESEARCHER** | 2.35 | 2.70 | 2.42 | 2.60 |
| **RESEARCHER-CLINICIAN** | 2.47 | 2.53 | 1.93 | 3.00 |
| **TECHNOLOGIST** | 2.43 | 3.43 | 1.86 | 2.29 |

The underlying data can be found in https://doi.org/10.25949/17009321.v1

Finally, we assessed the extent to which one’s own beliefs regarding the site of primary pathology are correlated with what one considers others’ beliefs to be in this regard. Out of the 101 participants, 98 selected a single location of highest importance (i.e., the survey allowed to indicate several damage locations as equally important). Of these 98 participants, those who selected IHC, OHC and SG as having the highest importance for Q1 also selected IHC, OHC and SG in Q2 (i.e., ‘what others believe’) respectively, see Table 2 for Spearman correlations analysis. Those who selected the SV as the highest important lesion site for Q1 didn’t think others maintained this “dogma” (i.e., no clear trend for any other damage site). This same trend was maintained for correlation analysis between Q1 and Q3 (i.e., excluding noise damage). If respondents thought the primary cause of age-related hearing loss was IHC, OHC or SG damage in Q1, they maintained this believe even excluding noise damage. Interestingly, only excluding noise damage analysis, people who believed SV was the main cause of presbycusis maintained this believe in Q3.

**Table 2.** Spearman correlation analysis between questions (Q1 vs. Q2 and Q1 vs. Q3) considering the primary pathology selected.

|  | Q2: IHC | Q2: OHC | Q2: SG | Q2: SV |
| --- | --- | --- | --- | --- |
| Q1: IHC | r=0.48, 95% CI [0.31 0.66] | r=-0.37, 95% CI [-0.55 -0.18] | r=-0.09, 95% CI [-0.29 0.11] | r=-0.02, 95% CI [-0.22 0.18] |
| Q1: OHC | r=0.05, 95% CI [-0.15 0.25] | r=0.51, 95% CI [0.33 0.68] | r=-0.47, 95% CI [-0.65 -0.30] | r=-0.18, 95% CI [-0.38 0.02] |
| Q1: SG | r=-0.32, 95% CI [-0.51 -0.13] | r=-0.23, 95% CI [-0.43 -0.04] | r=0.58, 95% CI [0.42 0.74] | r=0.11, 95% CI [-0.09 0.30] |
| Q1: SV | r=-0.31, 95% CI [-0.50 -0.12] | r=0.10, 95% CI [-0.10 0.30] | r=0.07, 95% CI [-0.13 0.27] | r=0.11, 95% CI [-0.09 0.31] |
|  | Q3: IHC | Q3: OHC | Q3: SG | Q3: SV |
| Q1: IHC | r=0.69, 95% CI [0.55 0.84] | r=-0.01, 95% CI [-0.21 0.19] | r=-0.18, 95% CI [-0.37 0.02] | r=-0.5, 95% CI [-0.67 -0.33] |
| Q1: OHC | r=-0.07, 95% CI [-0.27 0.12] | r=0.5, 95% CI [0.32 0.67] | r=-0.4, 95% CI [-0.58 -0.22] | r=0.08, 95% CI [-0.12 0.28] |
| Q1: SG | r=-0.22, 95% CI [-0.41 -0.02] | r=-0.16, 95% CI [-0.35 0.66] | r=0.64, 95% CI [0.48 0.79] | r=-0.18, 95% CI [-0.38 0.01] |
| Q1: SV | r=-0.49, 95% CI [-0.67 -0.32] | r=-0.33, 95% CI [-0.51 -0.14] | r=0.05, 95% CI [-0.15 0.25] | r=0.61, 95% CI [0.45 0.77] |

The underlying data can be found in https://doi.org/10.25949/17009321.v1

## Discussion

Contributions to hearing loss in general, and age-related hearing loss, or presbycusis, in particularly, likely arise from multiple sources and pathologies (Yu et al., 2021). Damage to outer hair cells and/or stria vascularis has a direct effect on hearing thresholds, and it is unsurprising these two mechanisms would feature in our respondents’ opinions as to the causes of presbycusis. However, our survey data suggest professionals in the field of hearing tend to consider damage to outer hair cells, rather than to stria vascularis (Wu et al., 2020), as the primary pathology in presbycusis. They also tend to believe other professionals in the field hold this opinion more strongly than they do, possibly because they rate their own knowledge of the indirect influence of any damage to stria vascularis on hearing thresholds (if this indeed is what they consider to be presbycusis) higher than the knowledge of others on this matter. Thus, according to our data, the dogma of the field is that damage to hair cells is the primary cause of presbycusis. Our findings raise the question as to why damage to stria vascularis might have been considered to be so by some researchers, given it is not, from our data at least, the view of most professionals in the field, nor is it the view they consider others to hold more strongly than they do themselves. One potential reason is the influence of critical thinkers on the field, particularly Schuknecht and Gacek (Schuknecht & Gacek, 1993), although there is little evidence to suggest that damage to stria vascularis was exclusively or primarily the cited cause of presbycusis in published studies, and such as view does not seem pervasive, given the outcomes of our study.

